# A coarse-grained model of disordered RNA for simulations of biomolecular condensates

**DOI:** 10.1101/2024.11.26.625489

**Authors:** Ikki Yasuda, Sören von Bülow, Giulio Tesei, Eiji Yamamoto, Kenji Yasuoka, Kresten Lindorff-Larsen

## Abstract

Protein-RNA condensates are involved in a range of cellular activities. Coarse-grained molecular models of intrinsically disordered proteins have been developed to shed light on and predict single-chain properties and phase separation. An RNA model compatible with such models for disordered proteins would enable the study of complex biomolecular mixtures involving RNA. Here, we present a sequence-independent coarse-grained, two-bead-per-nucleotide model of disordered, flexible RNA based on a hydropathy scale. We parameterize the model, which we term CALVADOS-RNA, using a combination of bottom-up and top-down approaches to reproduce local RNA geometry and intramolecular interactions based on atomistic simulations and in vitro experiments. The model semi-quantitatively captures several aspects of RNA-RNA and RNA-protein interactions. We examined RNA-RNA interactions by comparing calculated and experimental virial coefficients, and non-specific RNA-protein interaction by studying reentrant phase behavior of protein-RNA mixtures. We demonstrate the utility of the model by simulating the formation of mixed condensates consisting of the disordered region of MED1 and RNA chains, and the selective partitioning of disordered regions from transcription factors into these, and compare the results to experiments. Despite the simplicity of our model we show that it captures several key aspects of protein-RNA interactions and may therefore be used as a baseline model to study several aspects of the biophysics and biology of protein-RNA condensates.

## 1 Introduction

Biomolecular condensates formed from protein and RNA includes various types of membraneless organelles, having important and diverse roles in cellular activities.^1–5^ Although many mechanistic and biophysical aspects of these condensates remain incompletely understood, potential functional roles include ribosomal assembly in the nucleolus, protection of untranslated mRNA in stress granules, and RNA degradation in P-bodies. While subsets of specific proteins are thought to play central roles in the formation of these condensates, many other types of proteins and nucleic acids are also contained within them. Understanding and rationalizing the properties of condensates with multiple components would enhance our knowledge of cellular activities and the neurodegenerative diseases associated to the biomolecular condensates. For example, RNA may modulate the propensity for amyloid fibril formation in hnRNPA1A.^6^ Condensation of RNA alone can happen in RNA repeats, potentially leading to neurotoxicity. ^7,8^ Furthermore, protein–RNA condensates can selectively recruit other proteins. For example, proteins with charge-mixed domains are partitioned into nuclear speckles,^9^ and intrinsically disordered regions (IDRs) of transcription regulators with charge blocks are recruited into condensates formed by MED 1 IDR and RNA.^10^

Phase separation (PS) of protein and RNA is determined by both associative and repulsive interactions between proteins and RNA chains. ^11,12^ IDRs may contain polar, charged and aromatic residues, among which weak multivalent interactions have been found to be a major driver of PS of these proteins.^13–15^ The addition of RNA introduces negative charge, promoting PS in certain RNA-binding proteins and coactivators, whereas larger amounts of RNA can also inhibit PS.^16–18^ IDRs and RNA are sometimes found to exhibit so-called reentrant phase behavior, where PS only occurs at some intermediate RNA concentration, but neither at low or high RNA concentration. Mechanistically, this can be understood—at least in part—when PS of positively charged proteins that do not undergo PS on their own is induced by adding negatively charged RNA chains; further RNA addition may then prevent PS by solubilizing the positively charged protein.^5,18–20^ In the other direction, protein condensates may perturb the conformational properties of RNA and DNA.^21,22^

Molecular simulations have proven very useful to provide a microscopic view of molecular conformation and dynamics of IDRs.^23^ Atomistic simulations can capture the conformation of a single chain proteins or nucleic acids, and the dynamics of solvent and counter ions in IDP condensates,^24–26^ but they are limited by the time-scales and system sizes that can be studied. All-atom or coarse-grained (CG) molecular simulations with an implicit solvent can be used to study a larger number of protein chains on a longer timescales, at the cost of reduced resolution and potentially accuracy. This trade-off between spatiotemporal reach and resolution is governed by the concrete scientific question. A wide range of medium-resolution CG models have been developed for proteins, DNA and RNA to capture effects of protein secondary structure formation or nucleic acid base-pairing and neighbouring base interactions.^27–37^ These models commonly employ three or more beads per residue.

Complementing these approaches, hydropathy scale (HPS) models have been developed to represent disordered proteins and RNA with one-bead-per-residue mapping. These models generally do not represent local secondary structure formation (unless modified^38^) but enable simulations that reach longer time scales and larger length scales, and thus are a popular choice to study biomolecular condensates. The models have bonded force terms such as bond and, in some cases, angle potentials, and non-bonded interactions such as electrostatic and short-ranged interactions.^39–45^ While details vary across models, the strength of short-ranged non-bonded interactions generally depend on the sizes of the beads (*σ*) and a set of ‘stickiness’ parameters (*λ*) that capture the tendency of amino acids to interact with each other relatively to the surrounding solvent.

Homogeneous RNA chains such as polyA and polyU are often used in *in-vitro* experiments to study PS,^19,46,47^ and these RNA sequences often lack specific, well-defined structures, i.e. they can be modeled as mostly disordered RNA.^48^ As such, CG models may be particularly well suited to represent polyA or polyU chains, or to capture other RNA-protein systems where non-specific structures and interactions are the most important. We and other have previously described top-down approaches to parameterize CG models for proteins ^42,49–51^ based on experimental data. For RNA, the sparsity of experimental data makes detailed such purely data-driven parameter optimization difficult. Based on experimental data from SAXS and FRET of specific chains, a homogeneous RNA model with local chain stiffness have been proposed.^45^ Chou et al. proposed an HPS model of RNA, which was tuned using SAXS data and experimentally obtained dissociation constants.^52^ Even in the single-site model, the secondary structures of RNA are produced by employing explicit formalization of canonical base pairs and the neutralized beads.^53–55^ However, these models assume neutralized or reduced-charge beads for the backbone chain, and it is unclear to what extent this captures charge-driven aspects of PS with proteins.

Here we set out to develop a CG model for disordered RNA that can be used to study single-chain behaviour and phase separation of RNA in the presence of specific proteins. Previously, we have developed a protein HPS model, CALVADOS, using a data-driven topdown approach. ^49^ The interaction parameters were derived to match single-chain properties from experimental SAXS and NMR in a data-driven manner.^42^ Because of the symmetry between intra- and inter-molecular interactions in homotypic PS, CALVADOS also captures several aspects of PS of IDRs^44^ and multi-domain proteins.^56^ Here, we develop the RNA model in a consistent approach to CALVADOS 2^44^ to study protein–RNA condensates. Our model is aimed to capture situations when the main interactions of disordered RNA involve electrostatic interactions from the sugar-phosphate backbone and non-specific hydrophobic interactions from the nucleobases. Based on this idea, we represent each nucleotide by two beads, striking a balance between speed and resolution. As detailed below, we use a combined bottom-up and top-down approach to parameterize bonded and non-bonded interactions. We test the model on RNA–RNA and protein–RNA interactions in multi-chain systems, and demonstrate its utility by studying the selective partitioning of transcription factors into a condensate formed the IDR of the MED1 coactivator and long RNA molecules.

## 2 Results

### 2.1 Two-bead-per-nucleotide RNA model

We have developed a two-bead-per-nucleotide model of single-stranded (ss) RNA, consisting of one backbone bead and one base bead (Fig. 1a). The backbone bead represents the sugarphosphate backbone and is centred at the site of the phosphate atom, with ribose considered part of the backbone bead. The base bead is placed on the nitrogen atom that bonds the nucleobase to the ribose. A similar two-bead-per-nucleotide has been proposed for doublestranded DNA.^57^ Our model is intended to mimic the behavior of disordered RNA chains that do not adopt persistent secondary and tertiary structures. The backbone and base beads are RNA-sequence independent in our model, both to strive for a simple as possible model and because sequence-dependent parameters did not markedly improve agreement to experimental data at this level of resolution.^45^ However, we note that different disordered RNA sequences may in some cases have quantitatively different properties both in single chain and phase separation behaviour,^20,48^ limiting the range of questions the model can be used to study.

**Figure 1:**
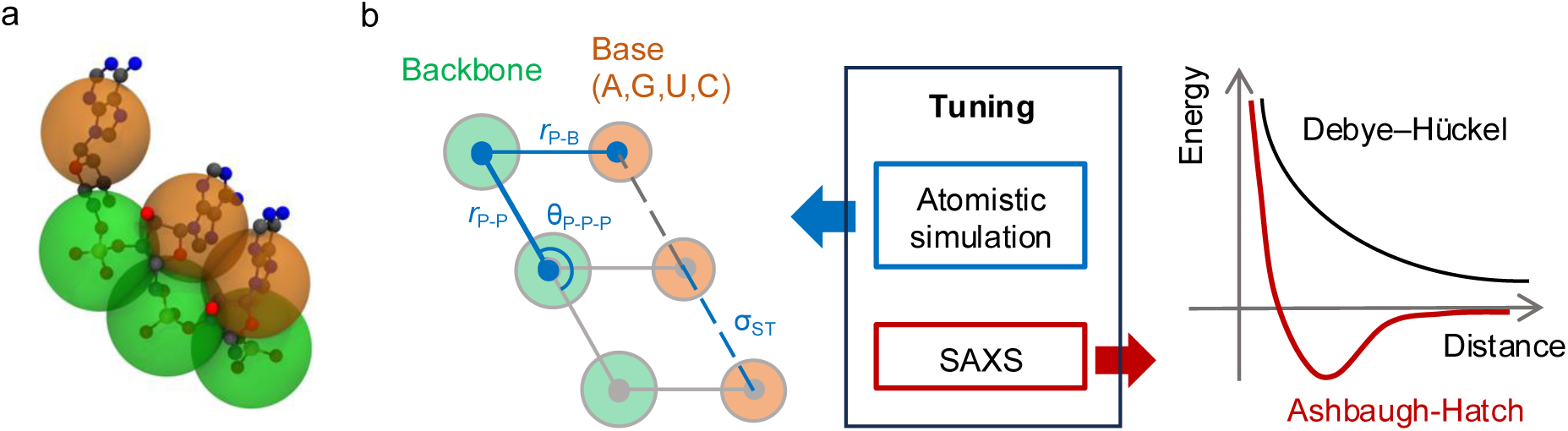
Two-beads-per-nucleotide model for single-stranded RNA. (a) RNA representation using the homogeneous coarse-grained (CG) model. The green and orange spheres represent the backbone bead and base bead, respectively. (b) We used atomistic simulation data to determine the model parameters for local strucural properties such as the backbone– backbone distance *r*_P_*_−_*_P_, backbone–base distance *r*_P_*_−_*_B_, backbone–backbone–backbone angle *θ*_P_*_−_*_P_*_−_*_P_, and the molecular size parameter for stacking interactions *σ*_ST_. Small-angle X-ray scattering (SAXS) data is used to determine the stickiness parameter of the Ashbaugh-Hatch potential.

### 2.2 Parameter tuning using a combined bottom-up and top-down approach

The potential energy function of the model includes bonded terms to describe bonds and angles, and non-bonded terms to capture short-range, electrostatic, and stacking terms (see Fig. 1b and Methods). After having defined the overall geometry of the model, the next step is to define the mass and size of the beads. The mass was computed as the sum of atomistic masses of the constituent atoms. We used previously calculated volumes that were based on the analysis of crystal structures^58^ to define the sizes of the coarse-grained beads by assuming Van der Waals spheres (Table 1). The sizes of the backbone and base beads are 0.69 nm and 0.62 nm, respectively, which are similar in size to the largest bead in the protein CALVADOS model (Trp; 0.68 nm).

**Table 1:** Parameters for beads used in this study (mass, molecular diameter and charge of beads, stickiness). We define a generic RNA nucleotide (R) as the arithmetic average of the four bases. The volumes are based on previously published data.^58^

| name | Mass [g/mol] | Volume [nm<sup>3</sup>] | $\sigma$ [nm] | charge [e] | $\lambda$ |
| --- | --- | --- | --- | --- | --- |
| Backbone RNA | 194.1 | 0.1761 | 0.6954 | -1 | 0 |
| Base RNA | 126.3 | - | 0.6238 | 0 | 1.18 |
| Base Ade | 134.1 | 0.1392 | 0.6430 | 0 | 1.18 |
| Base Uri | 111.1 | 0.1108 | 0.5958 | 0 | 1.18 |
| Base Gua | 110.0 | 0.1459 | 0.6033 | 0 | 1.18 |
| Base Cyt | 150.1 | 0.1150 | 0.6531 | 0 | 1.18 |

Subsequently, we determined force field parameters for local bonded terms such as bonds, angles and a local stacking potential using a bottom-up approach (Fig. 1b). The bond and angle potentials are modeled using a harmonic potential. The local stacking potential is represented by the Ashbaugh–Hatch (AH) potential and includes both a size parameter and a scaling parameter *n* that distinguishes between base-base interactions in neighbouring nucleotides from other base-base interactions (see Methods). We analysed previously published atomistic trajectories of a six-nucleotide sequence (UCCAAUC), ^59^ to extract the positions of the phosphorus atom and the nitrogen atom of the base bound to the ribose, and calculated the corresponding distributions of bonds, angles, and neighboring interactions to use as reference distributions. We initally set *λ_P_* = 0.2 and *λ_B_* = 1.2 for the non-bonded terms, and performed single-chain simulations of polyR30 (a 30-mer of a generic RNA) at a salt concentration of 100 mM and a temperature of 293 K for 200 ns. We manually scanned the parameters for the local force field terms to match the distributions from the CG model with those from the atomistic simulations. Since the microscopic properties are primarily influenced by bonded and stacking interactions, we fixed stickiness parameters *λ* at this step, and confirmed that the distributions did not significantly vary after tuning *λ*.

We validated that our model can accurately reproduce the distributions of microscopic properties. The equilibrium length and spring constant of the backbone–backbone bond are 0.59 nm and 1400 kJ/mol/nm^2^, respectively, while those of the backbone–base bond are 0.54 nm and 2200 kJ/mol/nm^2^. The equilibrium angle and spring constant are 3.141 rad and 4.2 kJ/mol/rad, respectively. The low spring constant for the backbone–backbone bond indicates the flexibility of the backbone chain. For the stacking parameters, the diameter is 0.4 nm and the scaling parameter is 15. These parameters produced bond and angle distributions that relatively closely matched those from the atomistic simulations (Figs. 2a– c). We note, however, that the base-backbone-backbone-base dihedral angle of the is not as well reproduced (Fig. 2d), as the A-form structure in UCCAAUC^59^ is not fully represented by our stacking function. Our model roughly reproduces location of the main peak, but it shows a long-tail distribution compared to the atomistic simulations (Figs. 2e and f). In addition to UCCAAU, we examined previously generated atomistic ensembles of polyA30 and polyU30.^60^ The bond and angle distributions do not significantly differ among these sequences, except for the dihedral angles, where adenine exhibits intense peaks compared to uracil (Fig. SS1). From these results, we conclude that our CG model successfully captures many key local structural properties of disordered RNA.

**Figure 2:**
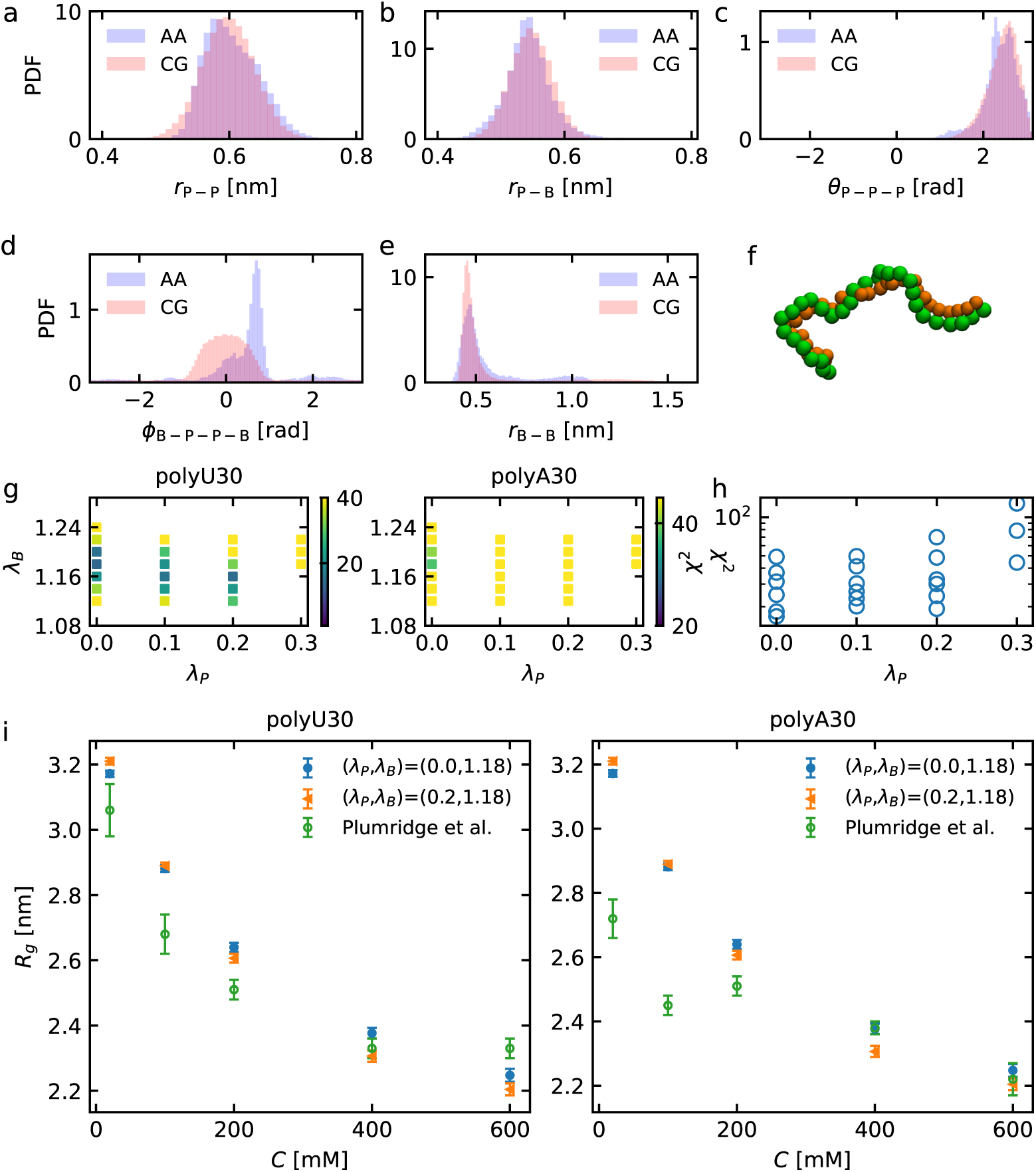
Parameterizing a coarse-grained RNA model. (a–f) We tuned a CG model of a 30-nucleotide RNA (polyR30) to mimic atomistic simulations of UCCAAUC ^59^ for the distribution of (a) the backbone–backbone distance *r*_P_*_−_*_P_, (b) the backbone–base distance *r*_P_*_−_*_B_, (c) the backbone–backbone–backbone angle *θ*_P_*_−_*_P_*_−_*_P_, (d) the dihedral angle of the base–backbone–backbone–base *ϕ*_B_*_−_*_P_*_−_*_P_*_−_*_B_ and (e) the base-base distance *r*_B_*_−_*_B_. (f) A representative structure from the single-chain simulation of polyR30 with backbone and base beads colored in green and orange, respectively. The results shown in (a–e) use the final values for the stickiness parameters (*λ*_P_ = 0 and *λ*_B_ = 1.18). (g–h) We parameterized the stickiness parameters of the Ashbaugh–Hatch potential in the RNA model to match the radius of gyrations *R*_g_ at varying ionic conditions, based on SAXS data of polyA30 and polyU30.^48^ (g) A grid search of *λ*_P_ and *λ*_B_. The reduced chi-square *χ*^2^ of *R*_g_ was calculated individually for polyA30 and polyU30, the color bar represents the *χ*^2^. (h) The sum of polyA30 and polyU30 data points was used to calculate the combined *χ*^2^. (i) *R*_g_ for polyU30 and polyA30 using selected parameters.

We then proceeded to determine the stickiness parameters for the RNA backbone, *λ*_P_, and for the base, *λ*_B_, using a top-down approach with the goal of reproducing the global dimensions of RNA chain. We utilized the radius of gyration (*R*_g_) of polyU30 and polyA30 at varying ionic concentrations (20–600 mM) based on SAXS measurements.^48^ Assuming that the backbone bead is hydrophilic and the base bead is hydrophobic, we performed a grid search over *λ*_P_ *<* 0.3 and *λ*_B_ *>* 1.0. Based on *χ*^2^ of *R*_g_ (see Methods), we find the optimal hydropathy parameters around *λ*_P_ = 0 and *λ*_B_ = 1.18 for both polyA30 and polyU30 (Fig. 2g) and when analysed together (Fig. 2h). At *λ* = 0, the AH potential describes purely repulsive interactions,^61^ i.e., no contribution from hydrophobic attractions. Since the radius of gyration does not change significantly for different values of the stickiness (Fig. 2i), we employed a simple scenario of purely hydrophilic–hydrophilic intentions for the backbone beads, i.e. *λ*_P_ = 0. For the base bead, *λ*_B_ = 1.18 was chosen. As point of comparison, *λ*(Glu) = 0 and *λ*(Trp) = 0.99 for the least and most sticky amino acids in CALVADOS 2, indicating that the RNA backbone is as non-sticky as glutamate and the nucleobases are on average more sticky than tryptophan. These values can also be compared to an earlier RNA HPS model based on atomistic charges that has a slightly negative stickiness,^39^ and our model is overall more sticky than that single-bead model. We note that these parameters produced semi-quantitative accuracy for *R*_g_ at varying ionic strengths when compared to polyU30, whereas the *R*_g_ of polyA30 at ionic strengths below 200 mM was not well captured (Fig. 2i).

We test the generality of our model across different sequence lengths by comparing apparent scaling exponents from our simulations with those calculated from SAXS measurements of ssDNA.^62^ Despite slight differences in the sugar between ssRNA and ssDNA, their properties as polymer chains—when disordered—are similar in FRET measurements.^63^ We performed additional single-chain simulations of polyR10–100 to calculate the scaling exponent. Our CG model shows scaling exponents of 0.66 at 225 mM and 0.56 at 525 mM (Fig. SS2). In SAXS measurements,^63^ 225 mM were used to derive a scaling exponent of 0.71 for polydA and 0.62 for polydT, while at 525 mM, the Flory exponent is 0.66 for polydA and 0.60 for polydT.^63^ Thus, the compaction in our model lies between that of polydA and polydT at 225 mM, whereas at 525 mM, our model is more compact than both.

### 2.3 Testing the accuracy of the model for intermolecular interactions

We evaluate the performance of our model, which is fine-tuned based on intra-molecular RNA interactions, to predict intermolecular RNA interactions by comparing calculations and experimental measurements of the second virial coefficient, *B*_2_. *B*_2_ is a measure of intermolecular interactions, where *B*_2_ *>* 0 indicates repulsive interactions and *B*_2_ *<* 0 indicates attractive interactions. In the context of molecular simulation, *B*_2_ has for example been used to assess protein–protein interactions in the development of CG models.^42,64,65^ We performed simulations of 400 polyR30 chains in cubic simulation boxes (Fig. 3a), and used the radial distribution function, *g*(*r*), for the center of mass of polyR30 to compute *B*_2_ (see Methods). *g*(*r*) *<* 1 at a distance *r* lower than 10 nm indicates repulsive intermolecular interactions, and *g*(*r*) increases as the ionic concentration rises, making the chains less repulsive (Fig. 3b). At 600 mM ionic strength, peaks at small distances (*r <* 2 nm) indicates some RNA chains can come closer together, which was also visually observed in snapshots. *g*(*r*) converges to 1 at 15 nm across all ionic strength conditions. Similarly, the *B*_2_ values obtained from *g*(*r*) show decreasing repulsive intermolecular interactions in our model with increasing ionic strength (Fig. 3c). Compared to the SAXS measurements of polyU30 by Pollack et al.,^48^ our model relatively accurately predicts intermolecular interactions at ionic strengths above 200 mM. However, the repulsive interactions of these highly charged RNA chains are underestimated at lower ionic strengths.

**Figure 3:**
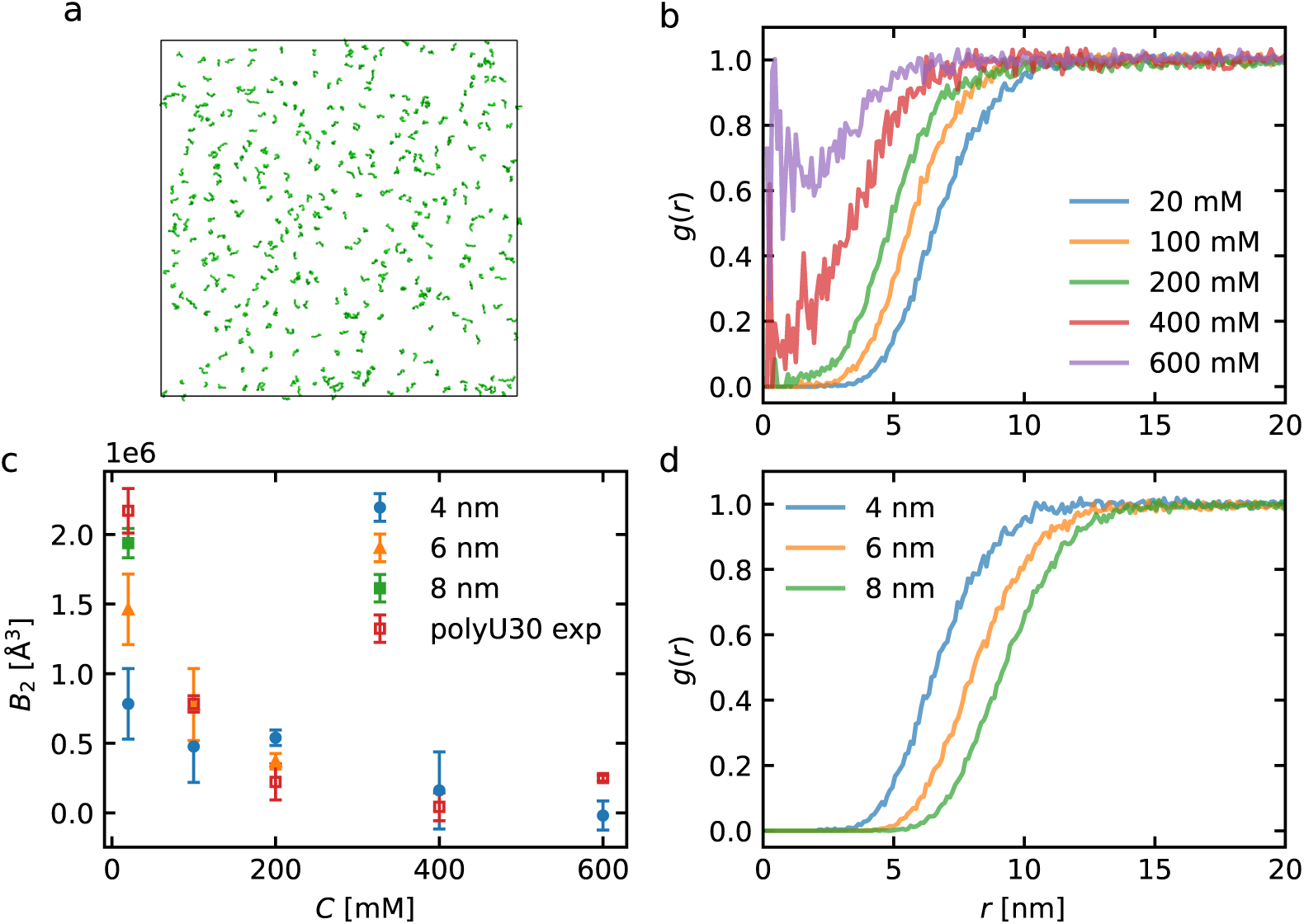
Prediction of intermolecular interaction of polyR30. (a) Snapshot of a simulation with 400 chains of polyR30. (b) Radial distribution function, *g*(*r*), at ionic strength 20– 600 mM. The cutoff distance of the Debye-Hückel (DH) potential, *r_c_*, is 4 nm. (c) Second virial coefficient, *B*_2_, calculated from simulations the different values of the *r_c_* for the DH potential. Experimental values for polyU30 by Plumridge et al.^48^ are shown for comparison. (d) Examining the effect of the cutoff distance on *g*(*r*) using *r_c_* =4, 6 and 8 nm (all at an ionic strength 20 mM).

We hypothesized that the discrepancy between our simulations and the experiments at low ionic strength at leats in part arises from longe-range electrostatic interactions beyond the cutoff used for the Debye-Hückel (DH) potential, as the DH potential decays slowly at low ionic strength. We therefore performed simulations using extended cutoff distances of 6 and 8 nm, in addition to the original 4 nm. Although increasing the cutoff distance resulted in lower values of *g*(*r*), *g*(*r*) from simulations with the longest cutoff distance (8 nm) converged at 20 nm (Fig. 3d). For *B*_2_, at the low ionic strength 20 mM, a longer cutoff distance of 8 nm reproduced the experimental values within the standard deviation error (Fig. 3c). At 100 mM, a 6 nm cutoff was sufficient to match the experiments within the standard deviation error. At higher ionic concentration (*>*200 mM), 4 nm was sufficient (Fig. 3c). Although the simulations with a 4 nm cutoff at 200 mM resulted in higher *B*_2_ values than the 6 nm cutoff, we speculate this discrepancy is due to uncertainty in the simulation and the procedures used to compute *B*_2_ (see Methods). Together, these results suggest that our model captures some, but not all aspects of the self-association of highly charge RNA molecules, and that long cutoffs are needed to model these charged systems at low ionic strengths.

### 2.4 Reentrant phase behavior of protein–RNA condensates

Following the parameterization and testing of our RNA model for intra- and intermolecular RNA interactions, we now assess its ability to describe protein-RNA condensation. We used the RNA model with the CALVADOS 2 model for IDRs, and calculate non-bonded protein-RNA interactions using combination rules (see Methods) to stay consistent with the definitions for intermolecular interactions between protein chains in CALVADOS 2. This is somewhat rationalized because the DH potential for the electrostatic interaction is identical for the IDR and RNA, and the short-range interactions are determined to balance the electrostatics.

We first studied the phase separation of intrinsically disordered RGG 3 domain of FUS (FUSRGG3) together with polyU40, where experiments have shown that FUSRGG3 phase separates together with intermediate concentrations of RNA, but not at low or high RNA concentrations.^47^ We hypothesized that the ability to capture such reentrant phase behavior would be a good test of the balance between hydrophobic and charge interactions, and inter-actions between RNA and protein, and note that other coarse-grained models have captured these effects.^36,39,52,66^ We used a fixed number of 500 protein chains (corresponding to a total protein concentration of 48.05 mg/mL) and performed multiple simulations with varying numbers of RNA chains at 293 K and 150 mM salt (Fig. 4a). In line with the experimental finding of reentrant phase transitions, we find that FUSRGG3 does not undergo PS in the absence of RNA. As also seen experimentally, FUSRGG3 undergoes PS in the presence of RNA, and these condensates become unstable at high RNA concentrations (Fig. 4b) Specifically, addition of RNA reduced the protein concentration in the dilute phase for up to 110 RNA chains, after which further RNA addition led to an increase in the protein saturation concentration (Fig. 4b). Stable droplets were not observed in systems with fewer than 20 RNA chains or more than 200 RNA chains.

**Figure 4:**
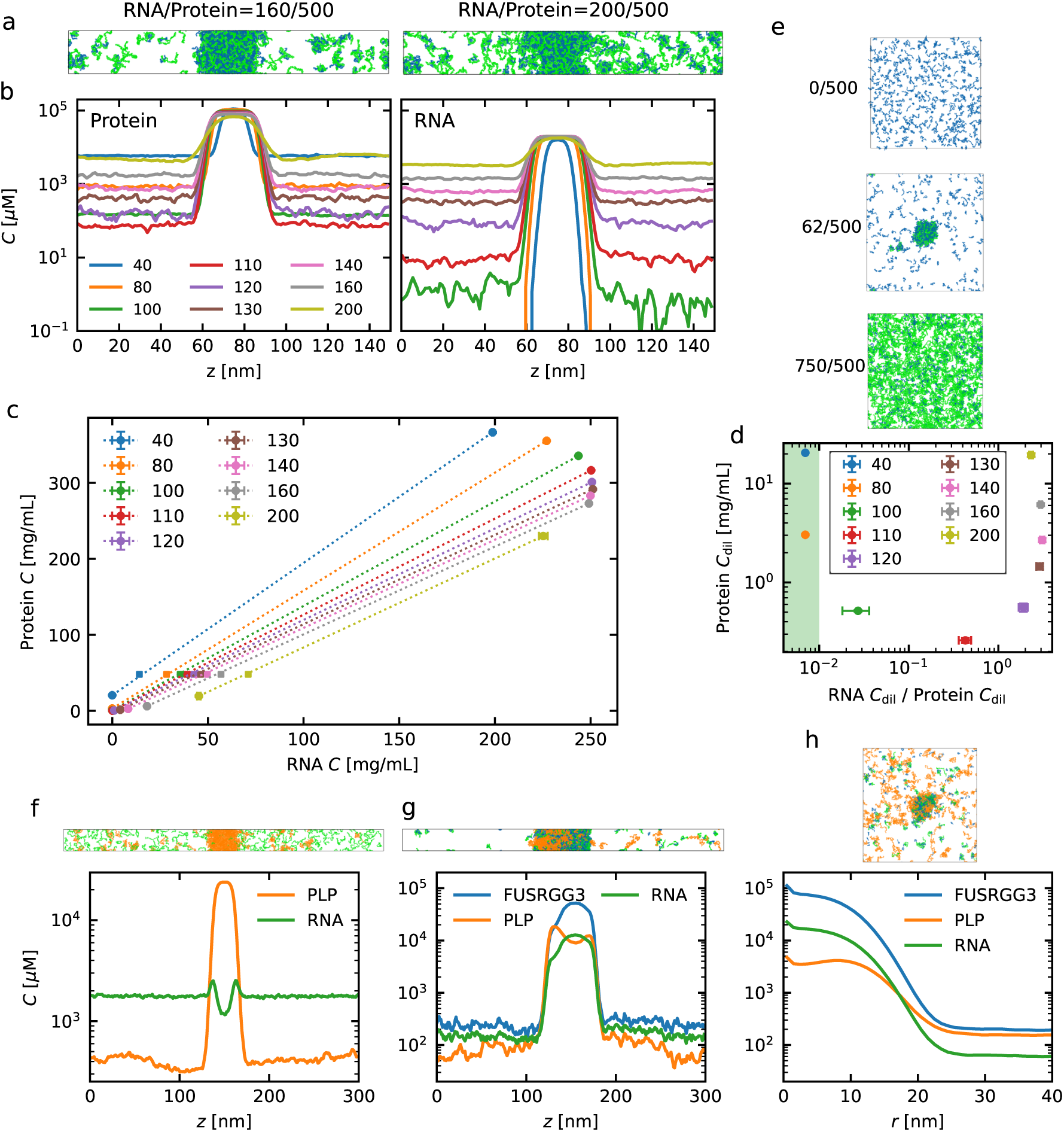
Slab simulations of protein (FUSRGG3) and RNA (polyU40). (a) Representative snapshots with protein colored in blue and RNA in green. The numbers of RNA and protein chains in the system are labeled above the simulation box. (b) Density profile of the molecular concentrations, *C*. The label shows the number of RNA chains. (c) Phase diagram with the concentration of RNA and protein at 293 K. Concentrations of dilute and dense phases obtained from the density profiles. The average density of the simulation box is also plotted. (d) Phase diagram in the dilute phase region. The *x*-axis represents the dilute phase ratio of RNA/protein (mass concentration). The lower limit of the x-axis where data can be estimated accurately is set at 0.01, and the region below this limit is shown in green. (e) Validation of the phase diagram using simulations with a cubic unit cell. (f) Simulation of the prion-like protein (PLP) and polyU40 in slab geometry. Simulation of systems with FUSRGG3, polyU40 and PLP in (g) slab geometry and (h) a cubic box.

We quantified the reentrant phase behavior using a two-dimensional phase diagram of protein and RNA concentrations.^47^ The concentrations of dense and dilute phases were obtained from the density profile (see Fig. 4b and Methods), which was then used to construct the phase diagram (Fig. 4c). In such phase diagrams, the tie lines connect the dilute and dense phase concentrations of the two components, and the slope of these tie lines contain information about the interactions between the two components.^14,67,68^ Positive values of the slope indicate associative PS. From our simulations, we find the slope to range from 0.15 to 0.23 in molar concentrations and 0.57 to 0.86 in mass concentrations, indicating that FUSRGG3 and polyU40 undergo associative PS. Second, we assess the conditions under which the saturation concentration of FUSRGG3 is lowest by showing the phase diagram in the dilute phase as a function of the RNA/protein mass ratio (Fig. 4d). We observed the lowest saturation concentration of FUSRGG3 when using 110 RNA chains, where the dilute phase exhibited an RNA/protein molar ratio of 0.116 and a mass ratio of 0.428. Since the nominal charge of FUSRGG3 in our simulations is +7 and that of polyR40 is -40, the neutral RNA/protein molar ratio is 0.175, corresponding to a mass ratio of 0.646. Thus, the lowest saturation concentration of protein was achieved near the neutralization point, indicating that the associative nature is mostly charge driven. Finally, the phase diagram (Fig. 4d) indicates that PS does not occur at the RNA/protein mass ratio above 3 when protein concentration is low (*<* 10 mg/ml).

To examine whether these results are non-specific to the slab geometry, we also performed additional simulations in cubic boxes. We therefore selected three points with RNA/protein mass ratios of 0 (no PS), 0.45 (PS), and 5.54 (no PS) at a bulk protein concentration of 2.8 mg/mL. The occurrence of PS in the simulations with a cubic box is thus consistent with the phase diagram obtained from simulations in slab geometry (Fig. 4e). To examine the effect of cutoff distance for DH potential, we performed additional simulations and produced a phase diagram. We observed a slight increase in the saturation concentrations of both protein and RNA (Fig. SS3a). However, apart from these subtle differences, the phase diagram obtained using 6 nm cutoffs was very similar to that using 4 nm (Fig. SS3b–d).

We compare the phase diagram obtained from our simulation to that from the experiments by Kaur et al,^47^ which was produced by examining PS conditions using FUSRGG3 and longer RNA chains. Their phase diagram shows the lowest saturation condition at an RNA/protein mass ratio of approximately 0.5, with protein concentration around 3.5 × 10*^−^*^2^ mg/mL. Our simulation results thus match the RNA/protein mass ratio at which protein saturation concentration is lowest; however, our simulations overestimate the saturation concentration. Regarding the maximum RNA/protein mass ratio, the phase diagram by Kaur et al.^47^ indicates that PS can occur at the ratios greater than 3. These results suggest that our simulations underestimate the propensity for PS. We speculate that this discrepancy arises in part because the long RNA chains (*>* 1000 nucleotides) used in the experiments exhibit a higher propensity for PS, but differences could also be due to ion-specific effects not captured by the DH model for these more charged mixtures. We also note that calculating the values for the left branch of the phase diagram of Fig. 4d for 40 and 80 RNA chains is challenging, because simulation convergence would require considerably longer simulation times or larger system sizes. We represent the x-axis values for RNA chains of 40 and 80 as 0.007, but we note that these exact numbers are not reliable.

To further examine the accuracy of the protein–RNA interaction, we investigate whether our RNA model can reproduce the mixing and de-mixing behavior of the prion-like domain of FUS (PLP). First, we performed slab simulations of PLP systems, where the estimated dilute phase concentration is 132 ± 8 *µ*M at 293 K and 177 ± 11 *µ*M at 297 K (Fig. SS4). Next, we performed simulations of PLP and polyU40 systems. Due to its net negative charge of -2, PLP shows repulsive interactions with RNA, resulting in polyU40 being excluded from the PLP slab (Fig. 4f). RNA does thus not reduce the concentration in the dilute phase (412.9 ± 28.8 *µ*M, see Fig. SS4), which is consistent with the experimental observations by Kaur et al.^47^ Finally, we performed simulations of the FUSRGG3, PLP and polyU40 system to test whether FUSRGG preferentially undergoes PS with polyU40, as observed by Kaur et al.^47^ In the slab simulation, PLP condensates were observed adjacent to, but not mixed within, the FUSRGG3–polyU40 condensate (Fig. 4g). To examine this observation in a different geometry we also performed simulations in a cubic box. Here we find a condensed phase with the concentration peaks of FUSRGG3 and polyU40 were located at the center, while the concentration peak of PLP is shifted away from the center (Fig. 4h). These results shows FUSRGG3 preferentially undergoes PS with polyU40, with PLP condensates positioned at the surface of the FUSRGG3–RNA droplet, consistent with the experimental observations by Kaur et al.^47^

### 2.5 RNA concentration influences the formation and selective partitioning of transcriptional regulators

To illustrate the utility of the RNA and CALVADOS 2 models to study biological problems, we studied the interactions between the long IDR from MED1 in the mediator co-activator complex (for simplicity we here term this IDR as MED1), RNA and IDRs from transcriptional regulators. We mimicked the 477-nucleotide long Pou5f1 enhancer RNA-1, as studied by Henninger et al,^18^ with our model.

As MED1 is highly charged (+43) it does not phase separate on its own (Fig. 5a). The addition of the RNA chain, which carries a charge of −477, compensates for the total positive charge, and the system is neutralized at an RNA/MED1 molar ratio of 0.09. We observed stable condensates also at an RNA/MED1 ratio of 0.2, but PS did not occur at high RNA/MED1 ratio such as 0.5 (Fig. 5a). Subsequently, we quantified the reentrant phase behavior of MED1 and RNA condensates using a phase diagram. We performed additional simulations at RNA/MED1 molar ratios of 0.05–0.3 to obtain the density profile (Fig. SS5a), and used the concentrations of MED1 and RNA in the dilute and dense phases to construct the phase diagram. The phase diagram (Fig. 5b) shows that the saturation concentration of MED1 decreases at RNA/MED1 ratios below 0.25 and increases at ratios above 0.25. On other hand, the dense phase reaches the highest concentration of both MED1 and RNA between RNA/MED1 ratios of 0.1 and 0.2. Similarly to the FUSRGG3 and polyU40 systems, both the dense phase concentration and its volume affect the dilute phase concentration. We find that the calculated phase diagram captures several aspects of the experimental observations by Henninger et al.,^18^ except for the left branch of the dilute phase (Fig. SS5b).

**Figure 5:**
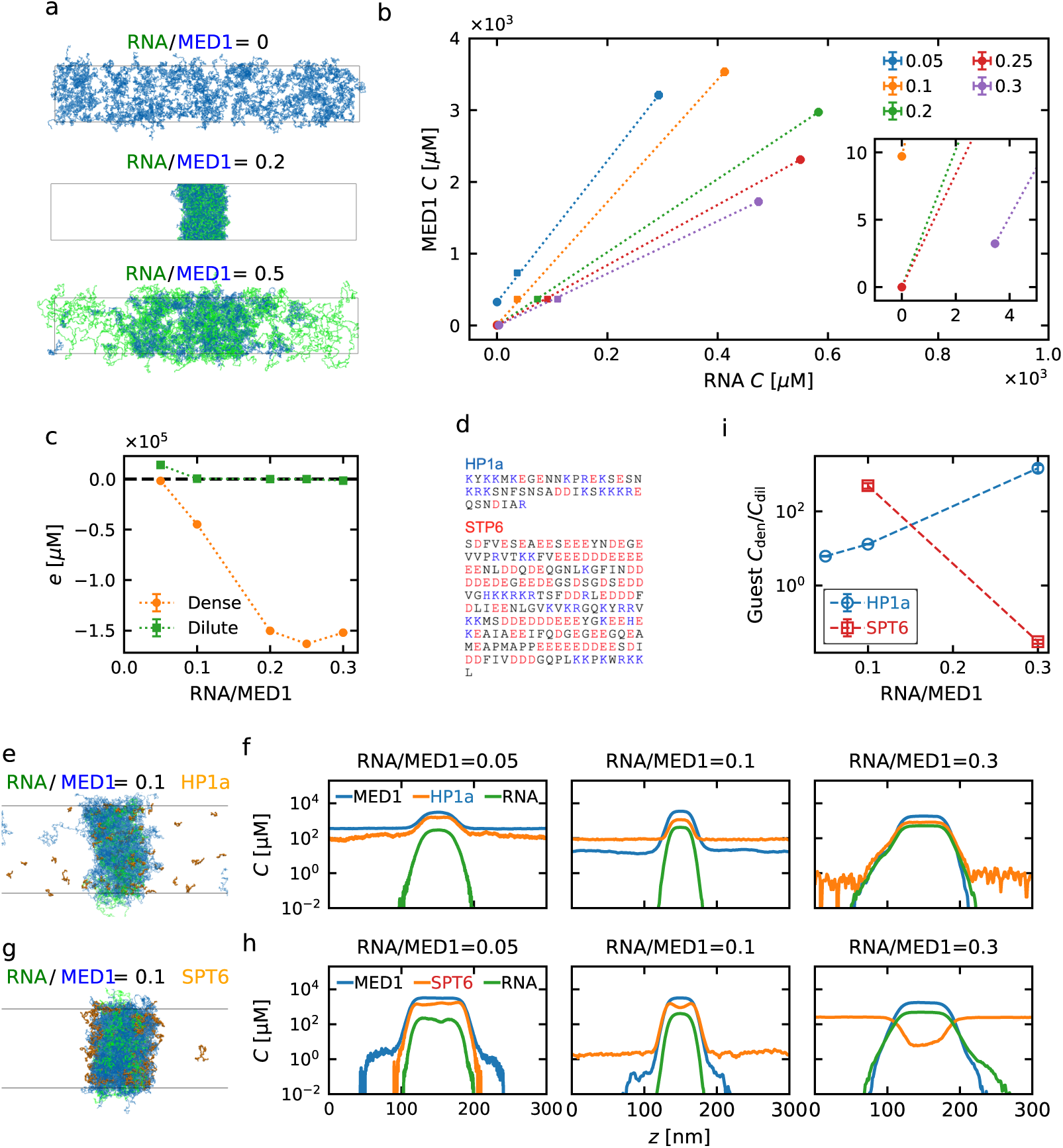
Reentrant phase behavior and selective partitioning of MED1–RNA condensates. (a) Representative snapshots of systems containing MED1 (blue) and RNA (green) with RNA/MED1 ratios 0, 0.2 and 0.5. (b) Phase diagram of the MED1 and RNA concentrations at 293 K, obtained from simulations with RNA/MED1 ratios of 0.05–0.3. The average density of the simulation box is also plotted with square markers. The inset highlights the region of the dilute phase. (c) Charge concentrations in the dilute and dense phases. (d) Sequences used for selective partitioning (HP1a and SPT6). Positively charged amino acids (R, K, H) are colored in blue, and negatively charged amino acids (D, E) are colored in red. (e, f) HP1a added to RNA–MED1 condensates. (e) A representative snapshot at an RNA/MED1 ratio of 0.1, and (f) the density profile at different RNA/MED1 ratios. (g, h) SPT6 added to RNA–MED1 condensates. (g) A representative snapshot at an RNA/MED1 ratio of 0.1, and (h) the density profile at different RNA/MED1 ratios. (i) Partitioning coefficients for HP1a and SPT6 at different RNA/MED1 ratios.

Since the phase diagram indicates that the composition of MED1-RNA condensates varies, we examined the changes in the electrostatic environments of the dilute and dense phase. We calculated a nominal macromolecular charge concentration as *Q*_MED1_*C*_MED1_ + *Q*_RNA_*C*_RNA_, where *Q* and *C* represent the nominal charge per chain and the molar concentration in a phase, respectively. The results shows that, at an RNA/MED1 ratio of 0.05, the dense phase is negatively charged (−1.76 × 10^3^ *µ*M), while the dilute phase is positively charged (1.40 × 10^5^ *µ*M) (Fig. 5c). An an RNA/MED1 ratio of 0.1, the dense phase is negative (−4.48 × 10^5^ *µ*M), and the dilute phase is nearly neutral (4.17 × 10^2^ *µ*M). At an RNA/MED1 ratio of 0.3, both the dense and dilute phase are negative with charge concentrations of −1.52 × 10^5^ *µ*M and −1.50 × 10^3^ *µ*M, respectively. These results indicates that the electrostatic environments of the phases are RNA-dependent. We note that these net charges refer only to the nominal charges of the macromolecules in our simulations, and in reality these would be partially compensated for example by small-molecule ions.

It has recently been shown that IDRs from different transcriptional regulators can partition into MED1–RNA condensates, and that the extent to which they do so depend on their charge and charge patterning.^10^ We therefore set out to study this selective partitioning using simulations. We further reasoned that changes in the electrostatic environment could alter the properties of MED1–RNA condensates, and therefore the extent to which different IDRs would partition into MED1–RNA condensates. To examine these questions, we performed additional simulations of MED1–RNA condensates with the inclusion of IDRs from transcription factors. We used the IDRs of heterochromatin protein 1a (HP1a) and SPT6 (Fig. 5d), as SPT6 has previously been shown to partition into MED1–RNA condensates, while HP1a does not.^10^ We performed partition simulations of HP1a and SPT6 (separately) using MED1–RNA condensates prepared at three different RNA/protein ratios (0.05, 0.1, 0.3). At the two lowest RNA/protein ratios we find that SPT6 partitions much more strongly than HP1a into the MED1–RNA condensates (Figs. 5e–h). We also find that the differential partitioning is highly dependent on the RNA/protein ratio so that at a RNA/protein ratio of 0.3 the situation reverses and HP1a now partitions much more strongly than SPT6, as quantified by the ratio of the concentration in the dense and dilute phases (Fig. 5i). These simulations correspond to points on a three-dimensional phase diagram (of MED1, RNA and SPT6/HP1a) and show, as expected, that the extent to which two components interact depends on the concentration of the third component. Our results also demonstrate that the rules for selective partitioning of transcriptional activators into for example coactivator–RNA condensates depend on the composition of these condensates.

## 3 Discussion

We have developed a two-bead-per-nucleotide model for disordered single-stranded RNA that is consistent with the CALVADOS 2 HPS model for IDPs.^42,44^ We term this model CALVADOS-RNA and the current version and parameter set as v. 1beta. We tuned the model parameters using a combined bottom-up and top-down approach, utilizing previously generated and publicly available atomistic simulation and experimental data. We showed that our model can capture several aspects of intermolecular RNA–RNA interactions and PS with IDRs using simple combining rules.

We hypothesize that the CALVADOS-RNA model, despite increased complexity and computational cost, offers some advantages over single-bead-per-nucleotide models. First, the mass and size of each bead are comparable to those in one-bead-per-residue protein models, potentially resulting in comparable energy scales for RNA-RNA and RNA-protein interactions. Second, the two beads represent the chemical nature with a polar and charged backbone and a more apolar set of nucleobases; the interactions with the charged backbone are dominated by the DH term, whereas the nucleobases interact with non-ionic interactions. This resulting flexibility from separating these interactions may help us to fine-tune the degree of RNA compaction in response to varying ionic strength.

There are also several limitations of our model. First, the model only represent unstructured RNA where specific base-base interactions are not important. It might be possible to combine the model with harmonic constraints to study more structured RNAs in a similar spirit to what we have recently demonstrated for proteins,^56^ but we have not yet tested this systematically. Second, for similar reasons, the model does not capture effects of RNA sequence. Sequence-dependent tuning may provide the potential to represent precise RNA– amino acid interactions^20^ and base-base interactions that can drive PS of RNA.^7,53,69^ More experimental data on structures and PS from different RNA sequences will be necessary for further refinement. Third, while we have shown that the RNA model can be used together with the protein CALVADOS model, we stress that it does not capture specific interactions between RNA and proteins; such interactions would be important to incorporate to study the role of domain structures in phase separation. ^70^ Fourth, we observed that the cutoff distance of the DH potential affects intermolecular interactions at low ionic strength. Although we show this effect for RNA-RNA interactions using virial coefficient (Fig. 2c) and RNA-protein interactions using the phase diagram (Fig. SS3), more systematic investigations are required. We speculate that it may be beneficial to adjust the cutoff distance based on the ionic strength. Fifth, despite the speed of the CALVADOS models, sampling low saturation concentrations (*<* 1 *µ*M) was not feasible in molecular dynamics simulations. Estimating thermodynamic parameters for systems that phase separate very strongly might require development and application of enhanced sampling methods.

Despite these limitations, we envisage that our RNA model will be useful to study the formation of protein–RNA condensates and their biochemical functions. Our simulations of MED1 condensates suggest that RNA concentration not only influences condensation but also alters the selectivity for IDRs of transcriptional regulators. Since MED1 forms transcriptional condensates at super-enhancer regions along with DNA sequences, RNA polymerase, coactivators and transcription factors,^71–73^ our results further suggest that the amount of RNA production may be one of the origins of dynamically fluctuating transcription rates in transcriptional condensates.^74^ Overall, the incorporation of RNA chains enables us to simulate membraneless organelles in a context more closely mimicking the cellular environment, allowing us to study the molecular mechanisms underlying their cellular activities and the diseases caused by their dysfunction.

## 4 Methods

### 4.1 Potential of two-beads-per-nucleotide model of RNA

The potential of the two-bead-per-nucleotide model consists of a bond term *U_b_*, an angle term *U_a_*, an electrostatic term *U*_DH_, a short-ranged AH term *U*_AH_, and neighbor stacking term *U*_ST_. The bonded potential is expressed as

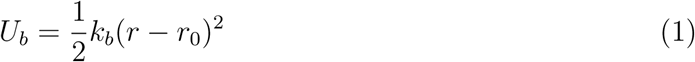

where the spring constant *k_b_* and equilibrium bond length *r*_0_ is determined for backbone– backbone distance and backbone–base distance, separately. The equilibrium length and spring constant of backbone–backbone bond are 0.59 nm and 1400 kJ/mol/nm^2^, and those of backbone–base bond are 0.54 nm and 2200 kJ/mol/nm^2^. The angle term is introduced to account for the stiffness of the backbone chain

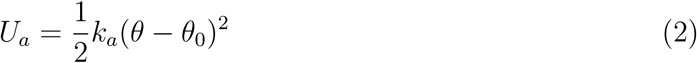

where the spring constant *k_a_* and equilibrium bond angle *θ*_0_ are determined for the backbone chain to be 4.2 kJ/mol/nm^2^ and 3.141 rad, respectively. The electrostatic interaction employs a DH potential,

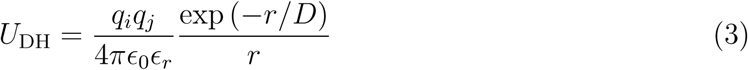

where *q* is the charge of the beads, i.e. -*e* for backbone bead and 0 for base, *ɛ*_0_ is the dielectric constant of water, *ɛ_r_* is empirical relationship for the temperature *T* dependency,^75^

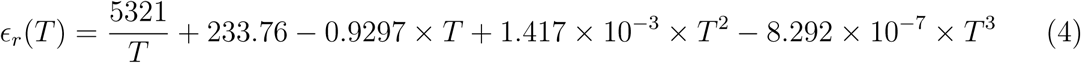

The Debye length is 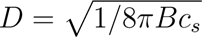 for Bjerrum length *B*(*ɛ_r_*) and ionic strength of monovalent ions *c_s_*. The DH potential is truncated and shifted at the cutoff distance of 4 nm unless stated otherwise. Short-range repulsive and attractive interactions are modeled by an AH potential,^61^

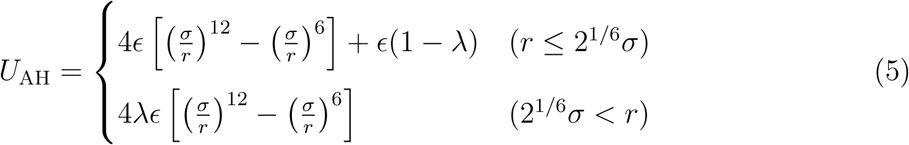

where *ɛ*=0.8368 kJ mol*^−^*^1^. For the molecular diameter and stickiness parameter, the combination rules, i.e., *σ* = (*σ_i_* + *σ_j_*)*/*2 and *λ* = (*λ_i_* + *λ_j_*)*/*2 are applied in the calculation of interaction between different types of beads. The AH potential is truncated and shifted at a cutoff distance of 2 nm. We express the stacking potential as

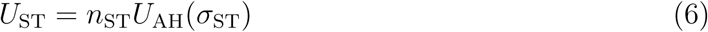

where *n*_ST_ is the scaling factor for the neighbouring attractive interaction and typically larger than 1, and *σ*_ST_ is smaller than the bead diameter for the intention to mimic anisotropic strong interaction between neighbours. We used *σ*_ST_=0.4 nm and *n*_ST_=15. *U*_ST_ was truncated and shifted at 2.0 nm as for the AH potential. We modeled neighbouring stacking interactions by an AH potential with a small *σ* instead of dihedral angle on base–backbone– backbone–base to maintain stable simulations at a large time step. We excluded the AH potential and electrostatic potentials for the pairs of beads with bond, angle, and neighboring stacking potentials.

We modeled all proteins with the CALVADOS 2 forcefield.^44^ We assume the above combination rules for *U*_AH_ for protein–RNA interaction.

### 4.2 Analysis of atomistic strctures

Atomistic structures of different RNA chains were obtained, specifically the UCCAAUC sequence from Bergonzo et al. generated by all-atom MD simulations,^59^ and the all-atom structures for polyA30 and polyU30, based on hierarchical chain growth methods, from Pietrek et al.^60^ In the analysis of these atomistic structures, bonds, angles, dihedrals, and base-base distances involving terminal residues were excluded.

### 4.3 Molecular dynamics simulation

All the simulations were preformed using OpenMM 8.1.^76^ Langevin integrator was used with time step 10 fs and friction coefficient 0.01 ps*^−^*^1^. Code and parameters for the CALVADOS-RNA v.1beta and protein CALVADOS models are available at https://github.com/KULL-Centre/CALVADOS.

In the single-chain simulation of polyR30, the initial configuration of polyR30 is placed at the center of a cubic box with a unit cell length of 30 nm. The conformation of polyR30 was an Archimedean spiral, where the backbone beads adopt this conformation in the *xy* plane, with the base aligned along the *z* axis, all oriented in the same direction. The temperature was set to 293 K, and multiple ionic concentrations *I* (*I*=20, 100, 200, 400, and 600 mM) were examined. Simulations were performed for 200 ns, with trajectories output every 100 ps. We disregarded the first 50 ns, and thus used the last 150 ns for analysis.

Multiple-chain simulations of polyR30 were performed using a cubic box with a unit cell length of 202.515 nm, into which 400 chains were randomly inserted. The concentration of polyR30 was 80.0 *µ*M.^48^ The conformation of polyR30 was the same Archimedean spiral as used in the single-chain simulation. The temperature was set to 293 K, and various ionic concentrations (*I* = 20, 100, 200, 400, and 600 mM) were examined. Additionally, the cutoff distance *r_c_* of DH potential was tested at 4, 6, and 8 nm, while the cutoff distance of AH potential was set to 2 nm.^44^ After energy minimization, simulations were performed for 10 *µ*s, with trajectories output every 1 ns. Disregarding the first 100 ns, the remaining frames were used for analysis.

The simulations of PS of FUSRGG3 and polyU40 were performed in a cuboidal box with unit cell dimensions [*L_x_*,*L_y_*,*L_z_*] = [20.0 nm, 20.0 nm, 150.0 nm]. A total of 500 chains of FUSRGG3 and 0–200 chains of polyR40 were inserted into the box, where the initial configurations consisted of fully elongated conformations aligned along the z-axis and centrally packed within the box. The temperature was set to 293 K, and the ionic strength was 150 mM.^47^ After energy minimization, simulations were performed for at least 4 *µ*s, with trajectories output every 10 ns. Disregarding the first 200 ns, the remaining frames were used for analysis. In the cubic simulations of FUSRGG3 and polyU40 condensates, the initial configurations and simulation procedures are the same as for those in the simulations of 400 chains of polyR30. The unit cell length was 100 nm. The number of FUSRGG3 chains were 500 and the number of polyU40 was 0, 62 and 750. After energy minimization, simulations were performed for at least 4 *µ*s, with trajectories output every 10 ns. Disregarding the first 1 *µ*s, the remaining frames were used for analysis. Slab simulations of PLP, PLP and polyU40, and FUSRGG3, polyU40, and PLP were conducted using the same initial configurations and simulation procedures as described in the slab simulations of the FUSRGG3 and polyU40 condensate. The number of chains for each component, slab geometry, simulation time, and equilibration time are detailed in Table S1.

The simulations of PS of MED1 and RNA condensates were performed in a cuboidal box with unit cell dimensions [*L_x_*, *L_y_*, *L_z_*] = [55.0 nm, 55.0 nm, 300.0 nm]. The RNA used was Pou5f1 enhancer RNA-1, a 477-nucleotide chain. The number of MED1 chains was set to 200, with 0–100 RNA chains. Since more than 10 RNA chains were necessary to form a sufficiently wide dense phase in this geometry, we instead prepared a system with 400 MED1 chains and 20 RNA chains (RNA/MED1 ratio 0.05). The initial configurations and simulation procedures were prepared similarly to the simulations of the FUSRGG3 and polyU40 system. The temperature was set to 293 K, and the ionic strength was 100 mM.^18^ After energy minimization, simulations were performed for at least 8 *µ*s, with trajectories output per 10 ns. Disregarding the first 1 *µ*s, the remaining frames were used for analysis. Similarly, slab simulations involving MED1, RNA, and guest LCDs (HP1a or SPT6) were performed. RNA/MED1 ratios of 0.05, 0.1, and 0.3 were produced using RNA/MED1 chain numbers of 20/400, 20/200, and 60/200, respectively. The guest IDRs (HP1a or SPT6) of 100 chains were inserted into the simulation box, except for the 0.1 RNA/MED1 ratio case, which used 200 chains of the guest IDRs. Simulations were run for at least 9.23 *µ*s and up to 21.59 *µ*s with trajectory output per 10 ns. The first 1 *µ*s were excluded from the analysis.

### 4.4 Analysis of the radius of gyration

We compared the radius of gyration of the simulation ensembles to experimental values to tune the stickiness parameters. Reduced chi-square *χ*^2^ was used to measure the fitness,

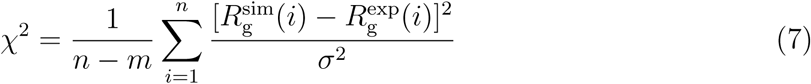

where *n* is the number of data points, *m* is the number of modeling parameters, that is two for the stickiness parameters of backbone and base bead, *R*^exp^ is the radius of gyration and *σ* is the standard deviation in the experiments. The data points contain polyU30 at ionic concentrations of 20, 100, 200, 400, and 600 mM, and polyA at 20, 200, 400, and 600 mM.^48^

### 4.5 Second virial coefficient

We quantified the intermolecular interactions between RNA–RNA based on the second virial coefficient *B*_2_,^42,65^

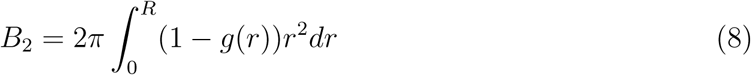

where *g*(*r*) is the radial distribution function of the center of mass of a single chain. In principle, *R* → ∞, but noise at high *r* significantly affects the result, particularly for repulsive systems. A correction for finite size effects was applied to calculate *g*(*r*).^42,77^ After checking the convergence of both *g*(*r*) and *B*_2_ with increasing *R*, we used *R* = 20 nm. The bin width of *g*(*r*) was set to 1 nm. The error bars for *B*_2_ were estimated by dividing the trajectories into 9 subsets of 1100 ns each, computing the radial distribution, and then calculating *B*_2_ for each subset. The standard deviation of *B*_2_ from these subsets was used to determine the error bars.

### 4.6 Concentration of dilute and dense phases

For the analysis of slab simulations, we centered the condensates along the long box axis and calculated density profiles at bin width 1 nm. In order to determine the position of the dense phase, dilute phase, and interface, we fitted the semi-profiles at *z <* 0 and 0 *< z* to the function,

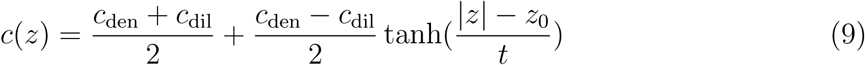

where *c*(*z*) is the density at position *z*, *c*_den_ and *c*_dil_ are the density of dense and dilute phases respectively, and *z*_0_ is the position to divide the phases and *t* is the thickness of the interface area. Then, the dense/dilute regions were defined using the cutoff parameter *β*,^44^ with |*z*| *< z*_0_ − *t* as the dense phase, and |*z*| *> z*_0_ + *βt* as the dilute phase. *β* was adjusted depending on the density profiles. For each time frame, we averaged the density of the two phases over the defined region to obtain average concentrations of the phases. We computed the mean value of the time-series values and estimated the standard errors using a time-blocking approach (https://github.com/fpesceKU/BLOCKING).

Concentrations were calculated from the density profile of each component as described above. In the MED1-RNA-Guest IDR systems, MED1 concentration profiles were used to determine the dense and dilute phase regions, and the concentrations of guest IDRs were then calculated for these regions.

## Data and code availability

The code and parameters for the CALVADOS-RNA v.1beta model are available at the CALVADOS GitHub page at https://github.com/KULL-Centre/CALVADOS. Additional data and scripts for this paper are available at https://github.com/KULL-Centre/_2024_ yasuda_RNA.

## Acknowledgements

I.Y. acknowledges support from JSPS KAKENHI (Grant JP23KJ1918) and, Keio University (Ishii-Ishibashi fund and Ushioda Memorial fund). S.v.B. acknowledges support by the European Molecular Biology Organisation through Postdoctoral Fellowship grant ALTF 810-2022. E.Y. acknowledges support from JST PRESTO (Grant JPMJPR22EE). The research was also supported by the PRISM (Protein Interactions and Stability in Medicine and Genomics) centre funded by the Novo Nordisk Foundation (NNF18OC0033950, to K.L.-L.).

## Competing Interests

K.L.-L. holds stock options in and is a consultant for Peptone. The remaining authors declare no competing interests.

## Supporting Information

**Figure S1:**
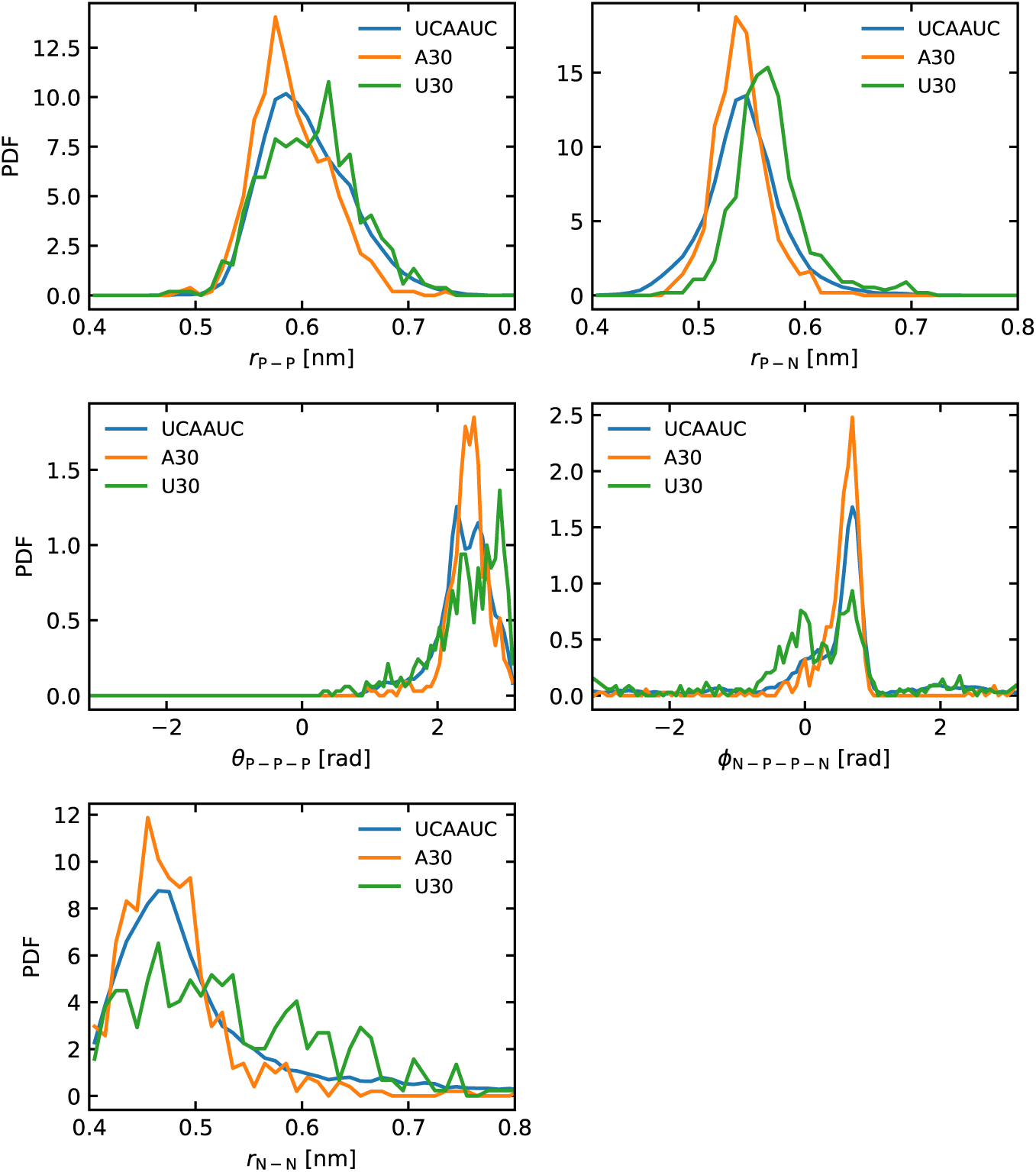
Comparison of atomistic structure ensembles of UCAAUC, polyA30, and polyU30 regarding bond, angle, and dihedral distributions. The structural ensembles were obtained from ref^59^ for UCAAUC, and from ref^60^ for polyA30 and polyU30.

**Figure S2:**
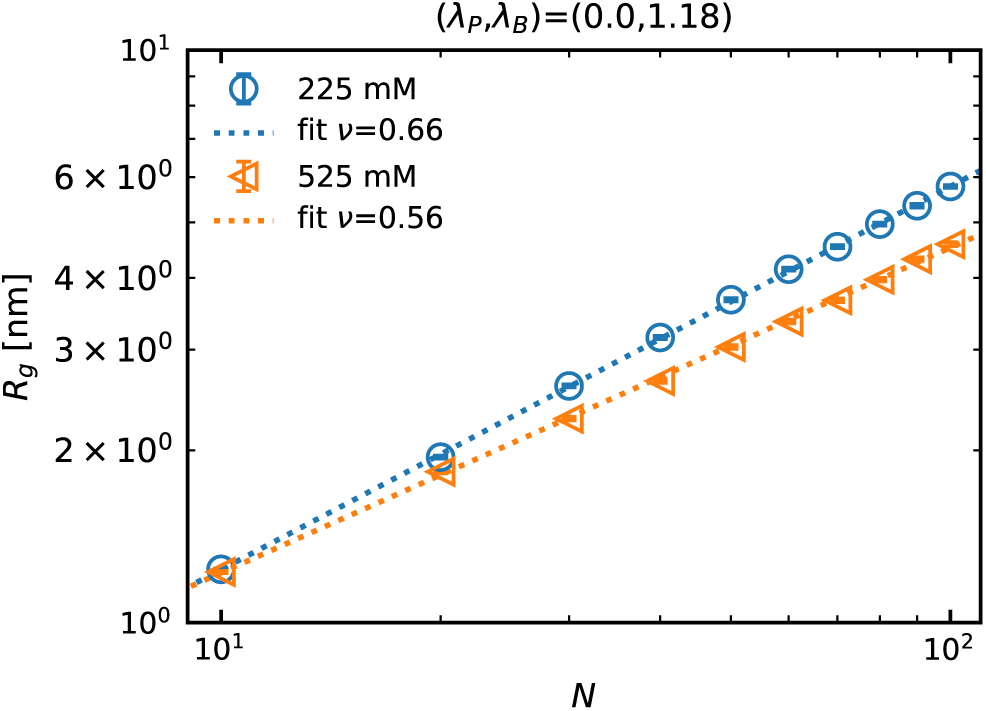
Radius of gyration *R*_g_ of polyR10–100 at ionic strength 225 and 525 mM. Fitting curve (dotted line) follows *R*_g_ = *R*_0_*N ^ν^*, where *R*_0_ and *ν* are fitting parameters and *N* is the sequence length.

**Figure S3:**
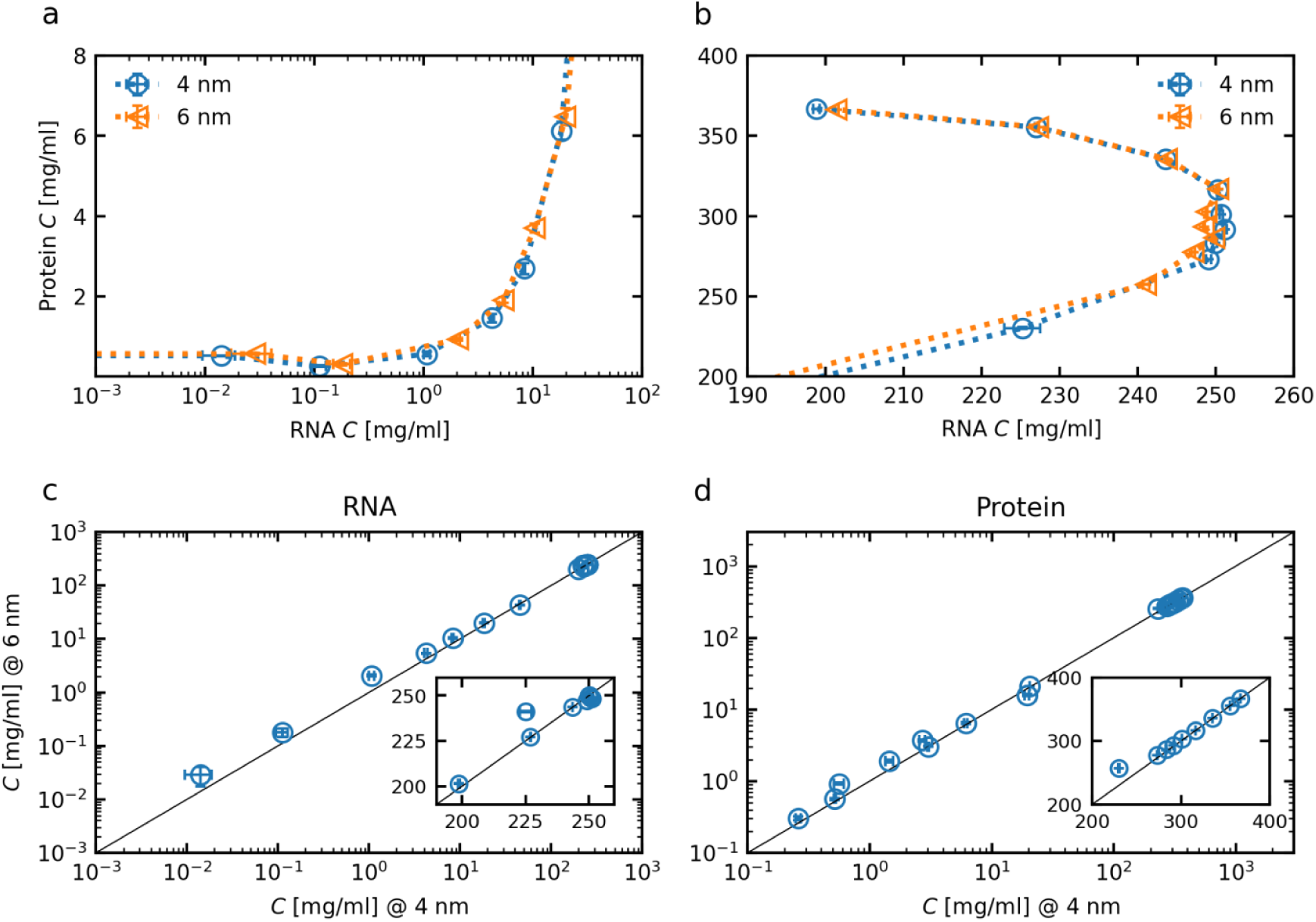
Phase diagram at cutoff distance of 4 and 6 nm for DH potential from simulation of FUSRGG and polyU40. (a) Dilute phase and (b) Dense phase. Comparison of concentration values for (c) RNA and (d) protein.

**Figure S4:**
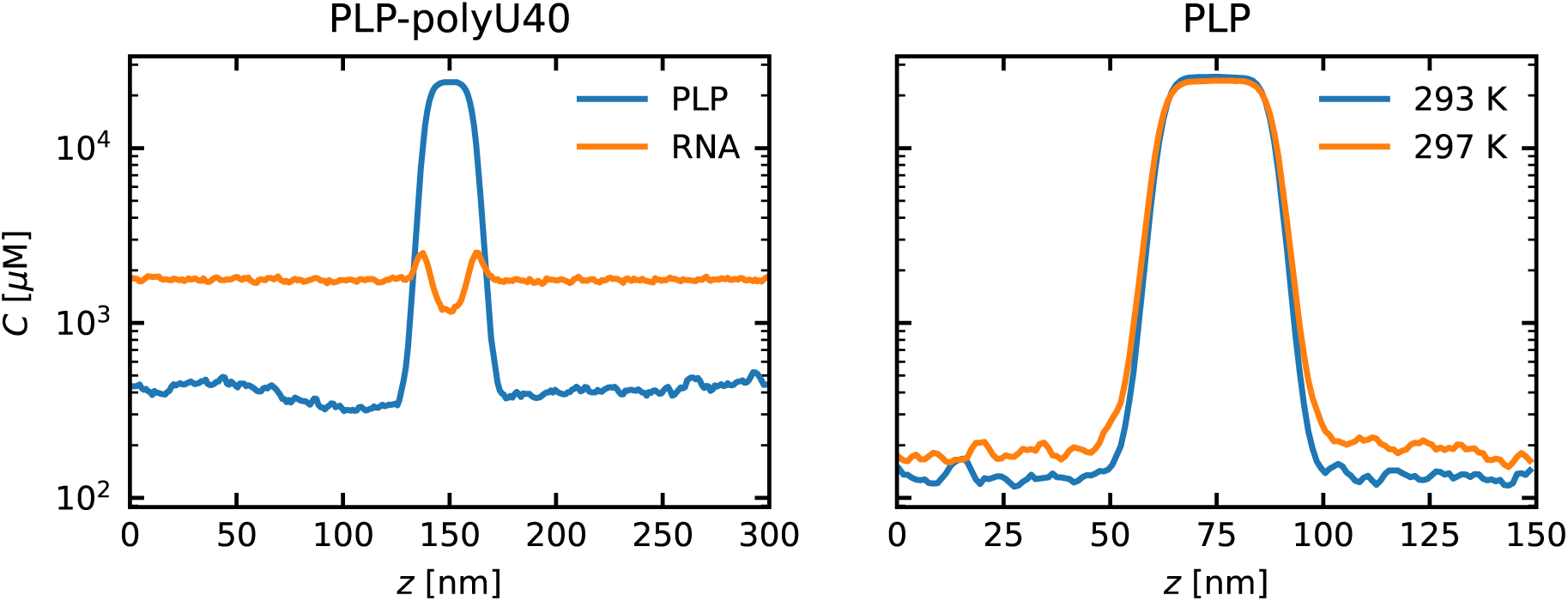
Density profile of PLP+polyU40 system at 293 K, and PLP system at 293 and 297 K.

**Figure S5:**
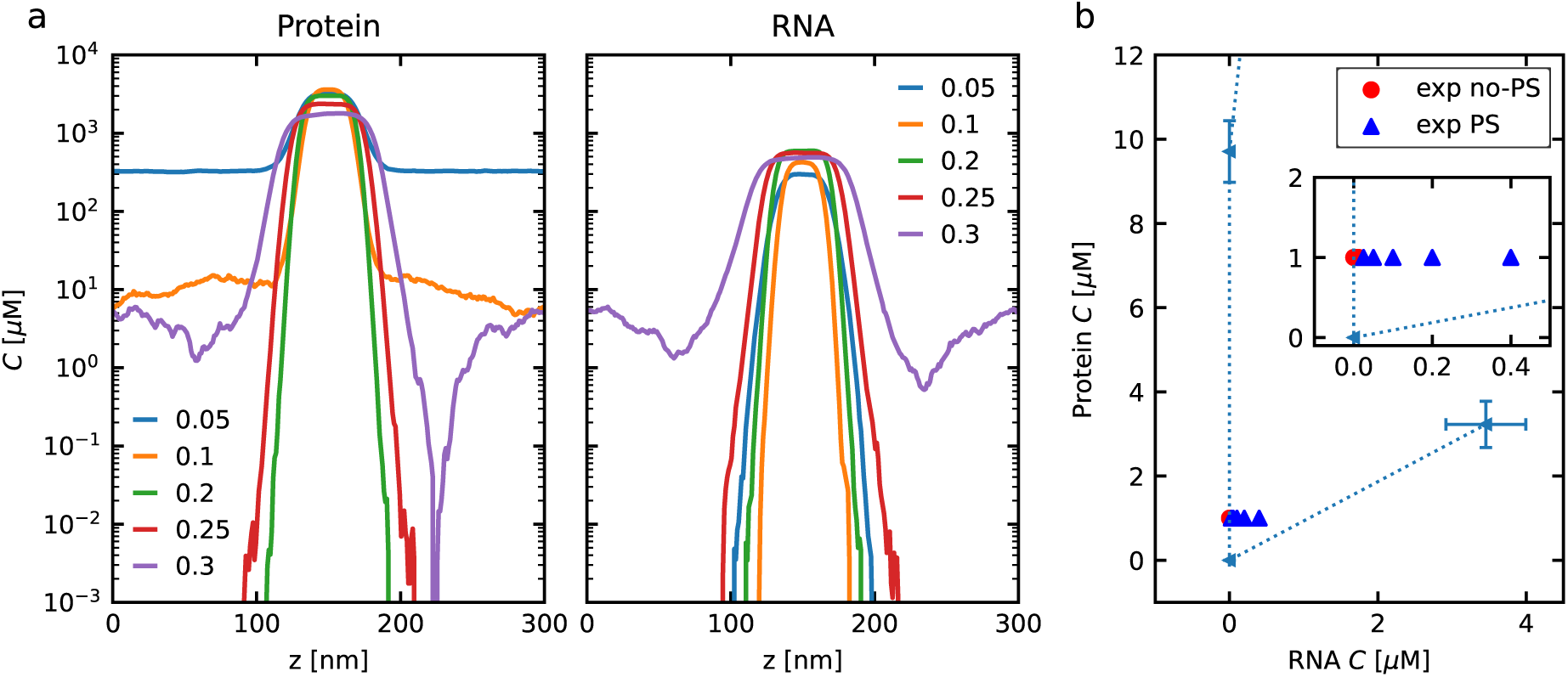
(a) Density profile of the MED1 and RNA system. (b) Phase diagram in the dilute phase. The binodal line from our simulation is compared with experimental observations of PS occurrence.^18^ The inset shows the region near the origin.

**Table S1:** Simulation setup for FUSRGG3, polyU40 and PLP systems. The table includes the number of chains for each component (FUSRGG3/PLP/polyU40), slab geometry [*L_x_*,*L_y_*,*L_z_*] in nm unit, simulation time *t*_run_, and equilibration time *t*_eq_.

| FUSRGG3/PLP/polyU40 | $[L_x, L_y, L_z]$ (nm) | $t_{\text{run}}$ ($\mu\text{s}$) | $t_{\text{eq}}$ ($\mu\text{s}$) |
| --- | --- | --- | --- |
| 0/328/0 | [20.0, 20.0, 300.0] | 12 | 3 |
| 0/328/256 | [20.0, 20.0, 300.0] | 12 | 3 |
| 473/164/128 | [20.0, 20.0, 300.0] | 4 | 1 |
| 473/164/128 | [110.8, 110.8, 110.8] | 15.4 | 1 |

## Notes

https://github.com/KULL-Centre/CALVADOS

https://github.com/KULL-Centre/_2024_yasuda_RNA

### Competing Interest Statement

KL-L holds stock options in and is a consultant for Peptone Ltd.

